# Divergent microbial structure still results in convergent microbial function during arrested anaerobic digestion of food waste at different hydraulic retention times

**DOI:** 10.1101/2022.11.21.517401

**Authors:** Minxi Jiang, Wendell Khunjar, Anjie Li, Kartik Chandran

## Abstract

In this study, two arrested anaerobic digestion bioreactors fed with food waste operated under different hydraulic retention times (HRTs) exhibited long-term stable volatile fatty acid (VFA) production performance including similar total yields (*p* = .085) with propionic acid (PA) being the most abundant VFA. Meta-omics analysis revealed distinct microbial structures (*p* = .02) at the two HRTs while there were no differences in potential and extant functionality as indicated by the whole-genome and whole-transcriptome sequencing, respectively. The highest potential (relative abundance of DNA sequence reads) and extant (relative abundance of mRNA sequence reads) functionality corresponded with PA production compared to other acids. The most abundant genus *Prevotella* produced PA mainly through the acryloyl-CoA pathway. Based on our results, the mechanistic basis for the similar VFA production performance observed under the HRTs tested lies in the community-level redundancy in convergent acidification functions and pathways, rather than trends in community structure.

## 1. Introduction

Arrested anaerobic digestion (AD) can be used for the synthesis of economically attractive volatile fatty acids (VFA) from diverse organic waste streams. VFA have a growing global market size of USD 13.41 billion by 2027 (Grand View Research, 2020). Compared to traditionally produced biogas from anaerobic digestion, VFA are more conducive to direct utilization in downstream applications such as bioplastic production (P. C. Lemos, 1998), biodiesel production (Vajpeyi & Chandran, 2015), and nitrogen removal in waste resource recovery facilities (WRRFs) (Guo et al., 2017).

Operational parameters such as HRT could be controlled to optimize the VFA yield and to manipulate the VFA composition before integrating into a stable downstream process (De Groof et al., 2021). Lim and coworkers reported an increased VFA yield in a mesophilic reactor running at pH 5.5 when HRT increased from 4 to 8 days (Lim et al., 2008). However, a further increase in HRT to 12 days didn’t result in an enhanced yield. This might be due to the potential COD loss to methane production under HRTs higher than 8 days (Yehuda Miron, 2000). As for the acid composition, acetic acid (AA), propionic acid (PA), butyric acid (BA), and valeric acid (VA) are the four most commonly produced acids, with the acid-speciation also being influenced by operating HRT. For example, a shift of the most abundant acid from AA to PA was observed when the HRT increased from 4 to 12 days (Lim et al., 2008) or 8 to 96 hours (Bengtsson et al., 2008). However, an unchanged AA-dominant production was also observed when HRT increased from 1 to 4 days (Dinsdale et al., 2000). Those variations of VFA composition as a function of arrested AD have a direct impact on downstream applications such as in the synthesis of polyhydroxyalkanoate (PHA) or when the VFA are used as carbon resources to enhance biological nutrient removal at waste resource recovery facilities (WRRFs). For example, when AA was the most abundant acid, the most produced PHA was PHB (Polyhydroxybutyrate) while PHV (Polyhydroxyvalerate) was produced in the majority when PA was the dominant carbon resource (P. C. Lemos, 1998; Pijuan et al., 2004). Besides, the AA-dominant product was reported to be preferred as the carbon resource for both denitrification and PHA accumulation as AA could directly enter the TCA (tricarboxylic acid) cycle for energy production (Elefsiniotis & Li, 2006; Vajpeyi & Chandran, 2015).

Several studies ascribed the variations of acid composition and yield to the microbial structural shift shaped by the operational HRT. The decreased AA fraction under HRTs higher than 5 to 8 days was correlated to the accumulated abundance of undesired aceticlastic methanogens under mesophilic conditions (Jankowska et al., 2015; Vanwonterghem et al., 2015). At 35°C and pH 5.9, an HRT of 10 days was reported to select *Olsenella* with the highest relative abundance and butyric acid was produced as the prevalent acid. However, when the HRT was decreased to 8 days, *Lactobacillus* exhibited the highest relative abundance producing lactic acid-dominated products (De Groof et al., 2021).Those results statistically correlated the VFA production performance with microbial structural abundance. However, other studies have suggested (without a detailed evaluation) that even with different microbial structures, similar VFA production could result (Chen et al., 2018). Besides, it was also found in AD plants that the microbial community composition shifted over time regardless of stable biogas production performance (Wu et al., 2016). Although these studies didn’t directly focus on the acidification functions, those results still indicated the potentially decoupled changes between performance, structures, and functions. Instead of correlating individual parameters, it is essential to have a complete insight into the nexus between the performance-structures-functions, especially for the dynamic changes in microbial structures and functions behind the long-time lumped acidification process of VFA production.

In this study, HRT 4 days and 8 days were selected to regulate the VFA production performance in laboratory-scale arrested anaerobic digesters from real food waste. It was hypothesized that under a longer HRT of 8 days (compared to 4 days), total VFA yield would be increased due to the extended reaction time, while AA might be produced as the dominant acid in both reactors considering the suppression of methanogens under selected operational conditions. Accordingly, we expected that the acidification bacteria would be enriched under longer HRT, and both the functional potential and activities of acidification would be enhanced.

To test these hypotheses, at each HRT tested, the VFA production performance was evaluated including both yield and composition. The structure of the microbial communities thus fostered was evaluated using single-gene (16S rRNA gene) amplicon sequencing and correlated to operational parameters. The global potential and extant function were compared based on the eggNOG database using whole genome and whole transcriptome sequence, respectively. Customized acidification metabolic networks were developed to describe different carbon transformation processes occurring during arrested AD and to reveal the specific changes in four acids (AA, PA, BA, VA) production pathways. Such approaches, aimed at opening the black box of anaerobic carbon transformation processes can be useful towards understanding the microbial structure-function-activity link and links-in turn to process performance, especially in complex engineered microbiomes such as arrested anaerobic digestion bioprocesses.

## 2. Methods

### 2.1. Reactors set-up and operation

Two 6L sequencing batch bioreactors with different HRTs (4 days and 8 days) were operated at 37.0±2.0 °C. Both reactors were inoculated with biomass from the anaerobic digestion process in Wards Island Wastewater Treatment Plant (New York, NY). The bioreactors’ pH was automatically controlled at 6.2±0.05 using 1M NaHCO_3_ (Feng et al., 2018; Jiang et al., 2013). The influent food waste was homogenized with a kitchen mixer and stored at -20 °C before feeding into the reactors. Both reactors were operated at a constant organic loading rate of 6.25 g total COD/L/day, which was within the practical range (3g COD/L/d to 15g COD/L/d) (Jiang et al., 2013; Magdalena et al., 2019).

### 2.2. Performance analysis

Bioreactor performance was evaluated based on duplicate measurements of chemical oxygen demand (COD) (Hach Chemical Co., Loveland, CO) and acidification metabolites such as lactic acid, formic acid, acetic acid, propionic acid, butyric acid, iso-butyric acid, valeric acid, iso-valeric acid, hexanoic acid, and heptanoic acid. The analyses of acidification metabolites were conducted through ion chromatography (IC) Dionex 2000 (Thermo Scientific, Sunnyvale, CA), which was equipped with an AS11 column (2 × 250 mm). The gas composition was analyzed by gas chromatography (Model 8610C, SRI instruments) with the column (Model ShinCarbon ST, Restek PA, USA) and thermal conductivity detector (TCD). The influent total solids (TS) and volatile solids (VS) were measured according to the standard method (APHA, 2005). The carbohydrate, protein, and lipid levels were quantified using the anthrone method (Morris, 1948), modified Lowry method (Thermo Fisher Scientific, IL, USA), and Folch method (Folch et al., 1957), respectively. All measured properties were summarized in SI Table 1. As defined in the SI Eq. (1) to Eq. (3), three parameters including VFA yield, methane yield, and individual acid fractions in total VFA were used to evaluate bioreactor performance.

**Table 1.**
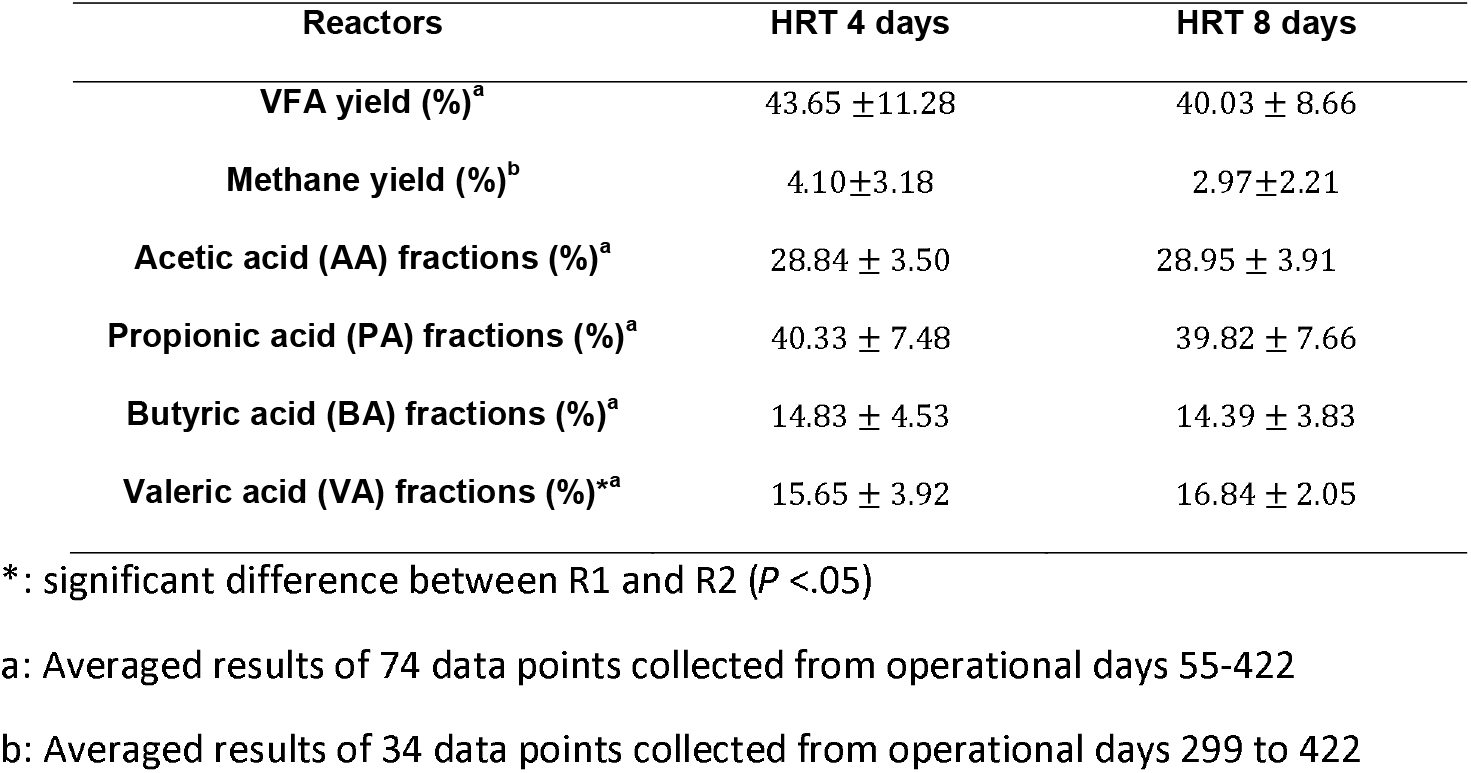
Overall performance results in the arrested AD bioreactors

### 2.3. Molecular analysis

#### 2.3.1. 16S rRNA gene sequencing and analysis

Biomass samples were collected weekly after the reactors reached a stable performance. DNA was extracted using DNeasy PowerSoil kit (Qiagen, MD, USA). Primer set 515F/806R targeting the V3-V4 region of both bacteria and archaea was used to amplify the 16S rRNA genes (Caporaso et al., 2011) and sequencing was performed on the Ion Torrent PGM platform (Thermo Fisher, MA, USA). The resulting sequences were processed using QIIME2 (2019.10). All the OTUs were identified at a 97% similarity. Silva.132 database was used as the taxonomic classification reference and results were illustrated using R (3.6.3) or python (3.8.0).

#### 2.3.2. Metagenome sequencing and analysis

The extracted weekly DNA samples were combined on an equal mass basis to reach a final weight of 500 ng DNA per HRT condition. The preparation of whole-genome libraries and sequencing were performed using Illumina HiSeq platform with pair-ended kits targeting 2×150 bp fragment length. The resulting pair-ended reads were screened with Trimmomatic (Phred score: 33, minimum length: 25 bp). Mothur was used to assemble the pair-ended reads into contigs, and sequences were aligned against the NCBI non-redundant protein database using Diamond Blastx (Identity percentage > 80%, maximum e-value: 0.00001). The global functional classification based on eggNOG was identified by MEGAN 6.0. Reads-based DNA percentages (as defined in SI Eq. (4)) were used to present the global functional potential.

#### 2.3.3. Metatranscriptome sequencing and analysis

Total RNA was extracted using the RNA PowerSoil Kit (MO BIO laboratories, Carlsbad, CA), and all extracted samples were combined on an equal weight (ng RNA) basis to reach a final weight of 500 ng RNA/condition. Library preparation and RNA sequencing were conducted using the Illumina HiSeq platform. The paired-end sequences were filtered, and adaptors were removed using the Trimmonatic command (Phred score: 33, minimum length: 70 bp). The filtered sequences were assembled using PEAR, and the ribosomal reads were removed with sortMeRNA. The coding regions were annotated by Diamond Blastp against NCBI non-redundant protein database (Identity percentage > 85%, maximum e-value:1e-15). The global functional classification based on eggNOG was further assigned by using MEGAN 6.0. Reads-based mRNA percentages were used to present global functional activities (as defined in SI Eq. (5)).

#### 2.3.4. Customized acidification metabolic networks and taxonomic origins

The reconstruction of the customized acidification metabolic networks of AA, PA, BA, and VA production was accomplished by integrating reference databases such as KEGG, IUBMB, as well as NCBI RefSeq. Genera presented in at least one sample with > 50000 RPM assigned and the species >10% in the genera were considered as a significant contribution to the total coding regions (CDS) of each system. Those enzymes identified in at least one sample and assigned to significant contribution species were selected in the customized metabolic networks. Each enzyme involved in the network was summarized with the corresponding gene names, gene length, and the involved reaction (SI Table 2). Based on the database, the graphs of the customized metabolic networks were drawn for each acid using Adobe Illustrator (2020). Given the uneven DNA potential of various genes, the transcriptional activity (mRNA RPKM as defined in SI Eq. (6)) of each gene was normalized by the genetic potential (DNA RPKM as defined in SI Eq. (7)) in the community to reveal the absolute transcriptional level with mRNA/DNA ratio (as defined in SI Eq. (8))(Xia et al., 2014). A customized Python script was used to specify the taxonomic origins of each DNA and mRNA read. Based on the DNA sequencing results (SI Table 3), the top five most abundant genera were selected, and their contributions to genes’ DNA RPKM (as defined in SI Eq. (9)) and mRNA RPKM (as defined in SI Eq. (10)) were calculated.

**Table 2.**
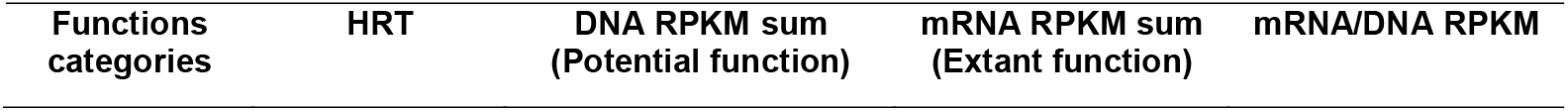
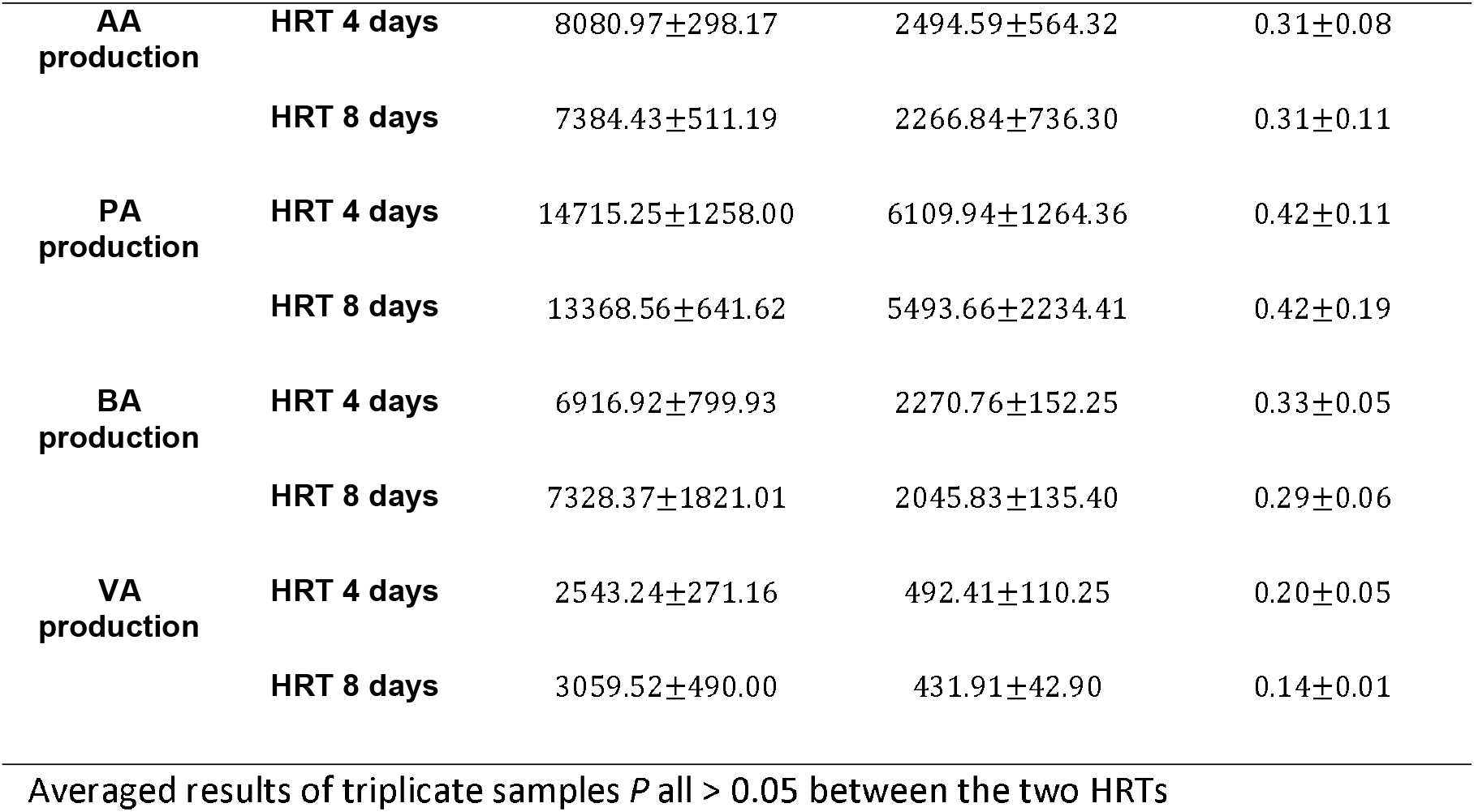
Potential and extant functions within the customized acidification metabolic networks

#### 2.3.5. Statistical analysis

Student t-tests were performed using the python package scipy.stats. and the statistical significance was considered with a confidence interval of 95% (*P* < .05). The similarity of microbial composition was estimated based on weighted-unifrac distance, and the db-RDA (distance-based redundancy analysis) was further conducted to explore the correlation between the formed microbial structure and operational factors (HRT and different food waste properties). The significant influence analysis of HRT and food waste properties on the microbial structure was estimated with an RDA permutation test using 999 permutations.

## 3. Results and discussion

### 3.1. Reactor performance

Overall, the VFA yield at the operating HRT of 4 days and 8 days was 43.65±11.28% and 40.03±8.66%, respectively (Table 1). These values were comparable to other reported yields from arrested AD reactors operated under similar conditions (35-37 C and pH 6-7). Vajpeyi found that the VFA yield was 38.7±12.9 % under HRT 2 days with food waste fermentation (Vajpeyi & Chandran, 2015). Feng observed the acidification yield from synthesized food waste was 44.3 % under HRT 4 days (Feng et al., 2018). In our study, increasing the HRT from 4 days to 8 days didn’t improve the VFA yield (*p* =.085), which was contrary to the hypothesis. Besides, both reactors were characterized by a negligible methane yield (HRT 4 days: 4.10±3.18%, HRT 8 days: 2.97±2.21%, Table 1, *p* = .35). This corresponded favorably to the previous work, which indicated a limited methane production at HRT lower than 5-8 days and 35-37°C (Kim et al., 2006; Vanwonterghem et al., 2015). Besides the potential limitation impact of HRT, the activities of methanogens might also be inhibited under HRT 8 days by the accumulated PA in the effluent (undissociated PA concentration: 0.34 ± 0.02 g COD/L based on pKa = 4.87), which was higher than the reported inhibition concentration (undissociated PA: 0.037 g COD/L based on pKa = 4.87) (Wang et al., 2009). Overall, the similar VFA and methane yields indicated that when methanogenesis was suppressed by the reduced HRT and high effluent PA concentration, the acidification community might be less sensitive to the HRT within the range of 4 days to 8 days.

As for the composition, PA was the most abundant acid regardless of the HRT (*p* = .71), which was also contrary to the hypothesis of AA-dominant products as widely reported in other studies. This PA-dominant product indicated the potential microbial contribution with organisms capable of PA production such as *Clostridia, Megasphaera, Prevotella*, and *Selenomonas* (Feng et al., 2018; Reichardt et al., 2014; Zhou et al., 2018). Alternatively, it could also be that metabolic pathways involved in PA production were possibly more actively expressed than other acid-production pathways. Meta-omics evidence for such possibilities is included in sections 3.2 and 3.4. The only difference in VFA production performance between the two HRTs was the significantly higher VA fractions under HRT 8 days (*p* = .035), which was not in alignment with the original hypotheses. Several studies have also reported increased VA fractions under extended HRT, which can be produced through chain-elongation by *Megasphaera* with ethanol and lactate being utilized as the electron donors (Contreras-Davila et al., 2020; De Groof et al., 2021; Kim et al., 2019; Wu et al., 2018). Therefore, VA production organisms such as *Megasphaera* might accumulate under HRT 8 days along with an enhanced VA production activity (Refer to sections 3.2 and 3.4).

### 3.2. Microbial ecology analysis from 16S DNA sequencing

#### 3.2.1. Bacterial and archaeal community structure

The Shannon diversity index was unchanged between the two HRTs (HRT 4 days: 3.60±0.56, HRT 8 days: 3.22±0.41, *p* = .06). *Bacteroidetes* phylum (HRT 4 days: 60.43 ±16.12 %, HRT 8 days: 51.62±12.53 %, *P* = .15) and *Prevotella* genus (HRT 4 days: 40.96 ±17.88 %, HRT 8 days: 40.23 ±11.71%, *P* = .91) were present at the highest relative abundance in both reactors (Fig.1). The *Bacteroidetes* phylum has been widely reported as the most abundant in anaerobic fermenters (Jia et al., 2013; Lim et al., 2014; Slezak et al., 2017; Zealand et al., 2018) and the human gut system (Ismail et al., 2011). The dominance of this phylum might stem from the tolerance to a mildly acidic environment and the wide-ranging hydrolytic capabilities to produce lipases, proteases, and carbohydrate-active enzymes from varying influents (Lim et al., 2014). At the genus level, *Prevotella* was reported to have the highest abundance in the gut system with a carbohydrate-rich diet (De Filippo et al., 2010), and was capable of producing PA from either acryloyl-CoA or methylmalonyl-CoA pathways (De Groof et al., 2021; Feng et al., 2018; Lim et al., 2014). The consistent prevalence of *Prevotella* in the two fermenters could likely contribute to the stable production of PA as the main acid product.

**Fig. 1.**
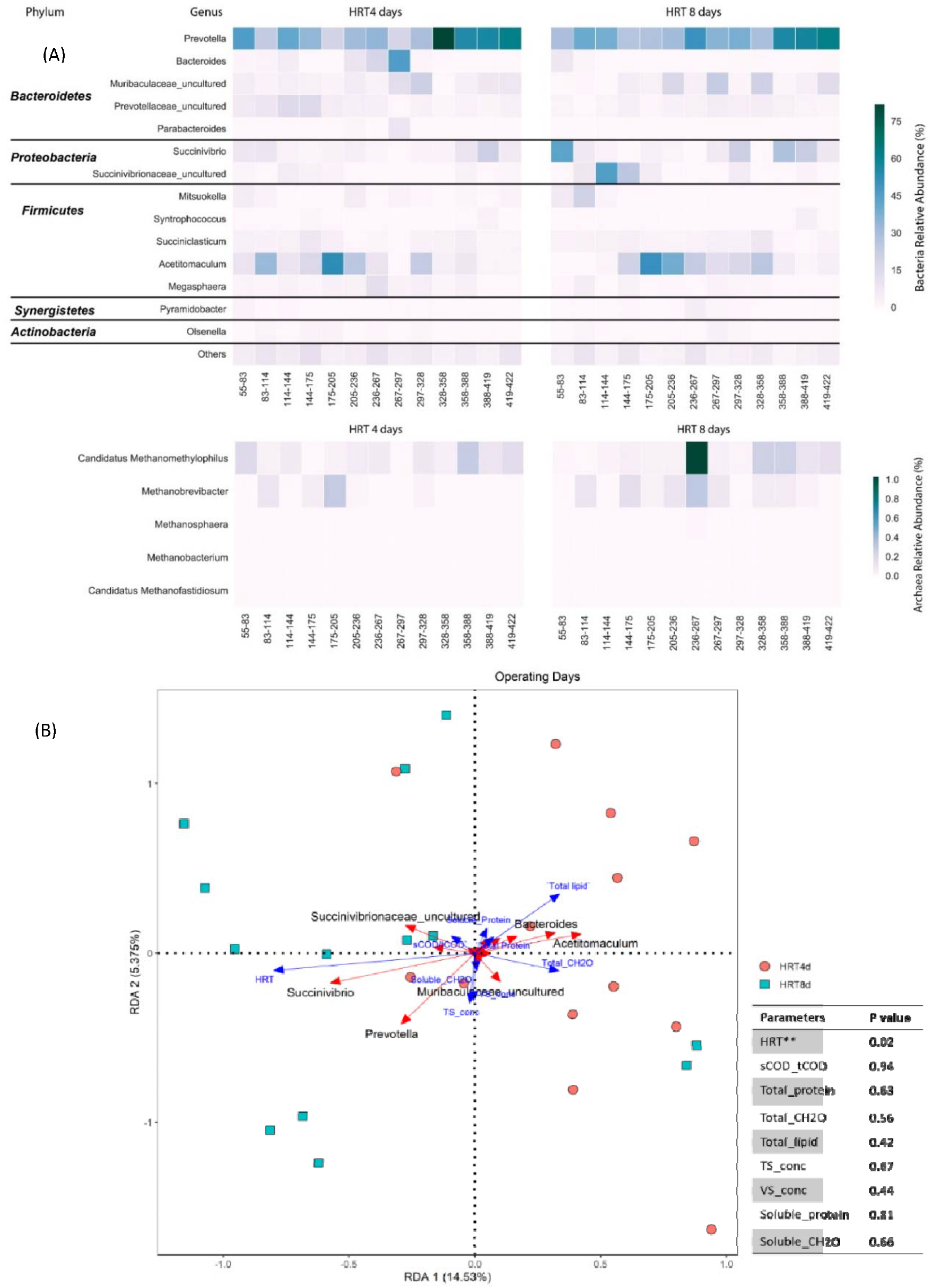
Microbial Ecology and significant impact of HRT and influent characteristics

Compared to bacteria, methanogenic archaea represented a minor structural component in both reactors (HRT 4 days: 0.14±0.09 %, HRT 8 days: 0.24±0.34 %, *p* = .32, Fig.1). This low abundance of archaea together with limited methane production demonstrated a successful suppression of methanogens. In both reactors, only methylotrophic methanogens (*Candidatus methanomethylophilus* HRT 4 days: 0.09 ± 0.08%, HRT 8 days: 0.17 ± 0.28%, *p* = .28) and hydrogenotrophic methanogens (*Metanobrevibacter* HRT 4 days: 0.04 ± 0.08%, HRT 8 days: 0.07 ± 0.09%, *p* = .33) were detected (Fig.1), while acetoclastic methanogens were absent under both HRTs (relative abundance = 0%, Fig.1). The hydrogenotrophic methanogens have been reported to coexist with acidification bacteria (Ziganshin et al., 2011) owing to a higher tolerance to the accumulated VFA and kinetics capable of overcoming lower HRTs (Magdalena et al., 2019). The exact reason for the presence of methylotrophic methanogens (although at low abundance) cannot be reconciled to their limited metabolic or kinetic capacities.

#### 3.2.2. HRT-shaped distinct flanking microbial ecology

Between the two HRTs, phylum *Proteobacteria* exhibited different abundance (*p* = .02). Besides, the variations of the abundance during the 16 different food waste batches cannot be neglected. The significance of controlled HRT and influent characteristics in shaping the overall formed microbial ecology was evaluated through the permutation test in redundancy analysis (RDA). The results indicated that HRT demonstrated a significant correlation to the variation of community-level ecology (*p* = .02, Fig.1). The higher HRT (8 days) was positively correlated to the relative abundance of *Prevotella, Succinivibrio*, and *Succinivibrionaceae_uncultured* (angle degree between arrows was < 90°, Fig.1), while negatively associated with the relative abundance of *Bacteroides, Acetitomaculum, and Muribaculaceae_uncultured* (angle degree between arrows was > 90°, Fig.1). This result was consistent with the hypothesis that different HRT would selectively facilitate the accumulation of certain acidification bacteria.

Based on biological principles, HRT selects the microbial community by their growth rate in continuously stirred tank reactors. If HRT < *θ*_*x min*_(minimal retention time for biomass), the bacteria would be removed faster than they are reproduced 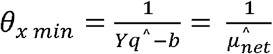, Y = growth yield, *q*^^^ = maximum substrate utilization rate, b = decay rate. 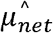 = maximum net growth rate) (McCarty, 2018). If the decay rate was not considered, the reported growth rate of *Prevotella* is *μ* 0.35/*h* (37°C) while for *Bacteroides* is *μ* 0.3 - 0.45/*h* (37°C) (Bacic & Smith, 2008; Ueki et al., 2007). A slightly higher growth rate of *Bacteroides* might benefit their reproduction under a lower HRT of 4 days, However, theoretically, the two selected HRT of 4 days and 8 days were enough for the growth of both bacteria, so the separation may not be mainly based on the growth rate.

On the other side, HRT also determined the reaction time inside the bioreactors. Considering that hydrolysis was the widely reported rate-limiting step during the acidification process, a longer HRT of 8 days may allow for more hydrolysis time, while a shorter HRT of 4 days may force the community to select species with efficient hydrolysis strategies such as a wider substrate range. As a result, the hydrolysis yield is similar between the two HRTs (HRT 4 days: 22.93 ± 8.31 %, HRT 8 days: 25.72 ± 9.20 %, *p* = 0.07). *Prevotella* and *Bacteroides* are known for their competitive capacity to decompose the hardly degradable polysaccharides with the produced polysaccharides utilization loci (PUL), especially for *Bacteroides*, which were reported with the highest number of experimentally verified PULs (Ausland et al., 2021). Besides, *Bacteroides* is also able to utilize proteins and lipids (De Filippo et al., 2010). In this study, it is likely that under HRT 8 days, *Prevotella* accumulated and gained energy for growth from more soluble molecular compounds released from hydrolysis, whereas under limited HRT like 4 days, *Bacteroides* was enriched to utilize slowly degradable polysaccharides as well as proteins and lipids.

From this result, it is concluded that HRT 4 days and 8 days might not mainly select the acidification community based on the growth rate. Instead, the capacity to efficiently utilize different substrates under limited reaction time differentiates the microbial structures.

### 3.3. Global-level functional potential and transcriptional activities

Although different microbial structures were shaped by HRT, no significant difference was detected in both the genetic potential (DNA reads-based alignment percentage) and transcriptional activities (mRNA reads-based alignment percentage) among all the 22 global functional classifications (*p* all > 0.05, SI Fig.1). This indicated that the distinctly formed acidification communities might redundantly encode and express convergent functions, as a result, generating similar VFA production performance.

In both reactors, the highest percentage of DNA reads was assigned to amino acid transport and metabolism (HRT 4 days: 11.47 ± 0.35 %, HRT 8 days: 10.76±0.10 %, *p* = .06, SI Fig.1). However, the highest percentage of aligned mRNA reads were assigned to carbohydrate metabolism in both reactors (HRT 4 days: 14.55±1.92 %, HRT 8 days: 12.10± 0.91 %, *p* = .15). This indicated a discrepancy between the functional potential and the transcriptional activities. Although the community might have the highest functional potential in amino acid utilization, carbohydrates could be actively transported and utilized as the main substrate and energy source during food waste fermentation. Compared to amino acid and carbohydrate metabolism, lipid metabolism was detected with less genetic potential (HRT 4 days: 2.69±0.08 %, HRT 8 days: 2.79±0.02 %, *p* = .15) and transcriptional activities (HRT 4 days: 2.77±0.39 %, HRT 8 days: 2.69±0.17 %, *p* = .78). This might stem from the adaptation of the microbial communities in the test bioreactors to a relatively small fraction of lipid contents in the food waste (< 3% w/w except for FW 144-175 days, SI Table 1), as well as the fact that lipids are hard to be enzymatically degraded due to the limited liquefication and potentially restricted mass transfer of lipase (Cirne et al., 2007).

### 3.4. Acidification metabolic networks

The sum of the DNA RPKM (HRT 4 days: 24952.88± 1724.88, HRT 8 days: 23804.51 ± 1207.99, *p* = .50) and mRNA RPKM (HRT 4 days mRNA RPKM: 9889.70±1687.46, HRT 8 days mRNA RPKM: 9011.00±2753.36, *p* = .07) of the total genes tracked within the acidification metabolic networks were unchanged between the two reactors. This resulted in the consistent mRNA/DNA RPKM ratio (HRT 4 days: 0.38±0.11, HRT 8 days: 0.40±0.09, *p* = .82). Those similarities in the meta-omics level in the acidification networks might be used as an indicator of the highly similar total VFA production performance.

#### 3.4.1 Acetic acid (AA) production

Recalling the performance results, there was no difference between the two reactors in terms of AA fractions (*p* = 0.29). Within the metabolic networks, seven pathways were reported for AA production, and all of them were found to be completely expressed in either HRT 4 days or 8 days (all genes involved in these pathways had mRNA/DNA RPKM >0, Fig.2). 7 genes involved in the last step of each pathway were used as the pathway indicators. It is noticed that *ackA* gene (pathway of acetyl-P to AA) displayed significantly higher DNA RPKM under HRT 8 days (*p* = .04, labeled in red text, Fig. 2). However, the mRNA/DNA ratio of gene *ackA* was not changed between the two HRTs. Besides gene *ackA*, other genes involved in the final steps of AA production pathways all didn’t exhibit any difference in terms of the mRNA/DNA ratio (*acs, aldH, pct, pauA, yccx, poxB, p* >0.05, represented in color black, Fig.2). This same mRNA/DNA ratio of all genes, as well as the same expressed pathways number, might contribute to the unchanged AA fractions between the two HRTs. In the meantime, it is suggested that instead of using only potential functional results from DNA, the mRNA/DNA ratio could exclude the unevenness from structural abundance and indicate the community-level activities, which in turn better correspond to the performance.

**Fig. 2.**
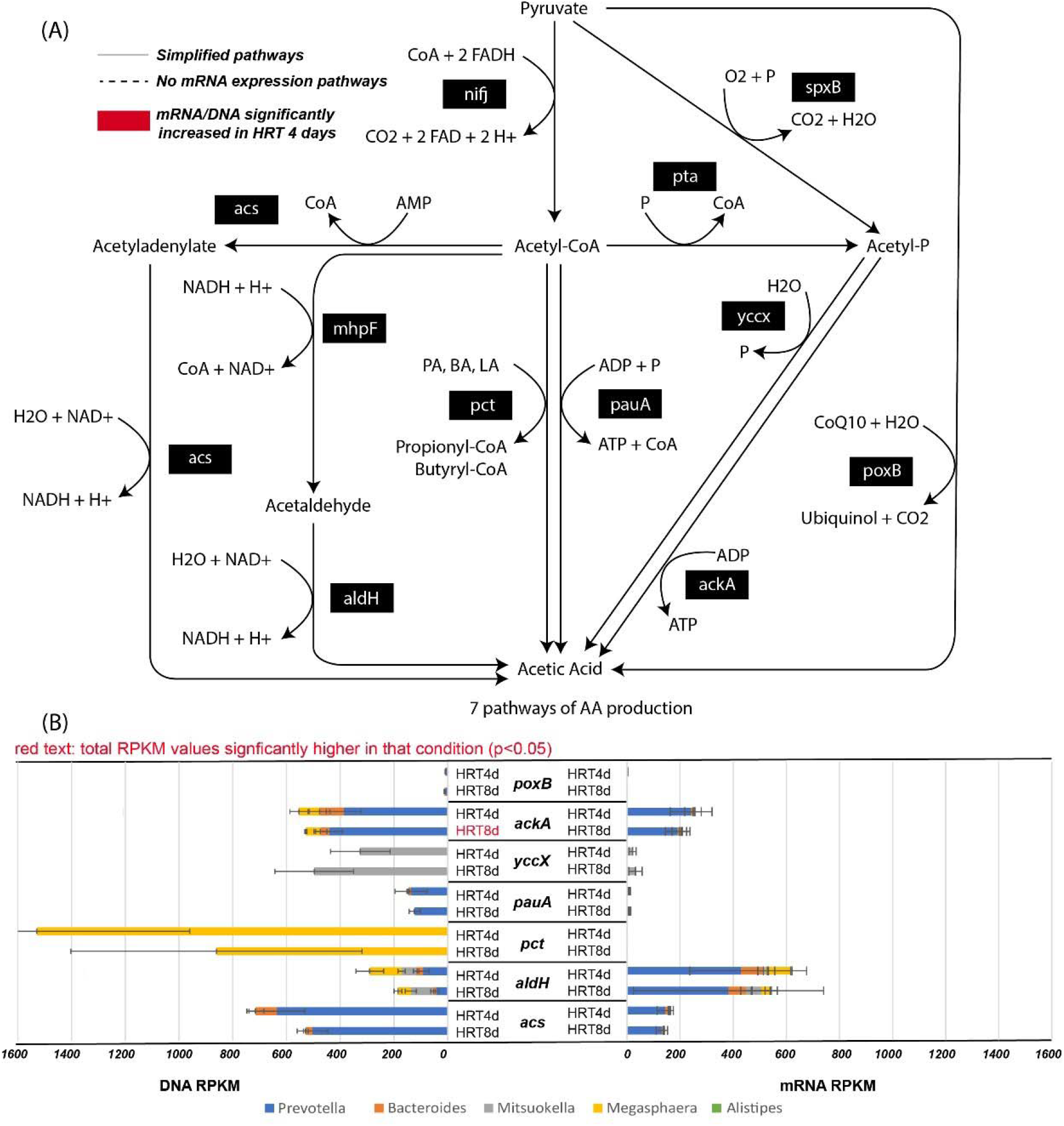
Acetic acid (AA) production metabolic networks and taxonomic origins

Different functional groups were found to produce AA via different pathways. *Prevotella* was capable to produce AA via four pathways in both reactors (Fig.2 stacked column left): (1) *acs* gene involved acetyladenylate to AA pathway (HRT 4 days DNA RPKM: 635.72±103.68, HRT 8 days DNA RPKM: 503.36 ±57.02, *p* = .14); (2) *ackA* gene involved acetyl-phosphate to AA pathway (HRT 4 days DNA RPKM: 386.53±65.43, HRT 8 days DNA RPKM: 441.70±50.01, *p* = .31); (3) *pauA* gene involved acetyl-CoA to AA pathway (HRT 4 days DNA PRKM: 136.54±59.64, HRT 8 days DNA PRKM: 122.64±20.64, *p* = .73); (4) *poxB* gene involved pyruvate directly to AA pathway (HRT 4 days DNA RPKM: 7.72±13.08, HRT 8 days DNA RPKM: 8.10±7.49, *p* = .97). *Megasphaera* could also generate AA from acetyl-CoA, but another gene *pct* encoded for the enzyme involved in this pathway (HRT 4 days DNA RPKM: 1529.09±568.93, HRT 8 days DNA RPKM: 860.76±541.64, *p* = .21). Similarly, the pathway from acetyl-phosphate to AA was also coded with another gene *yccx* in *Mitsuokella* (HRT 4 days DNA RPKM: 325.47±110.72, HRT 8 days DNA RPKM: 496.50±146.54, *p* = .19). These results suggested that the same AA production pathway was redundantly coded in different genera with distinct genes. Besides, the same gene might be widely distributed in many genera such as *aldH*. This gene encoded aldehyde dehydrogenase (NAD+), which catalyzed the reaction of acetaldehyde to AA (pathway 7). All five most abundant genera were coded with gene *aldH* (Fig.2 Stacked column left).This widely distributed genetic potential together with diverse pathways leading to AA production likely generate a community-level functional convergence even with changed microbial structures shaped by HRT. Although 7 potential pathways were redundantly coded in different genera to generate AA, only three pathways exhibited mRNA RPKM > 150 which include: (1) acetyl-P to AA (gene *ackA*), (2) acetaldehyde to AA (gene *aldH*), and (3) acetyladenylate to AA (gene *acs*). In all three pathways, *Prevotella* presented the highest mRNA contribution percentage in both reactors (*ackA*: HRT 4 days: 50.10±6.63%, HRT 8 days: 42.32±8.05%, *p* = .26, *aldH*: HRT 4 days: 45.07±7.06%, HRT 8 days: 45.73±16.79%, *p* = .95; *acs*: HRT 4 days: 84.72±3.03%, HRT 8 days: 86.74±6.19%, *p* = .65). These results indicated that during the arrested anaerobic digestion of food waste, AA is mainly produced by *Prevotella* through three pathways, which could be tracked by genes *ackA, aldH* and *acs*.

#### 3.4.2. Propionic acid production

Recalling the performance results, PA was always present as the most abundant acid in both reactors (HRT 4 days: 40.33±7.48 %, HRT 8 days: 39.82±7.66 %, *p* = .71). Compared to the other three acids, PA production also exhibited the highest DNA RPKM sum (HRT 4 days: 14715.25±1258.00, HRT 8 days: 13368.56±641.62, *p* = .17, Table 2) and mRNA RPKM sum (HRT 4 days: 6109.94±1264.36, HRT 8 days: 5493.66±2234.41, *p* = .70, Table 2) of all genes involved in the PA metabolic networks in both reactors-indicating the highest *potential* and *extant* functionality at the conditions studied. Within the PA metabolic networks (Fig.3), propionyl-CoA served as the important precursor and three pathways were merged at propionyl-CoA production before it was further converted to PA: (1) 1,2-propanediol pathway; (2) acrylate pathway, and (3) methylmalonyl-CoA pathway (Gonzalez-Garcia et al., 2017). In both reactors, only the methylmalonyl-CoA pathway was detected with all involved genes expressed (mRNA/DNA >0, no dash lines within the pathway, Fig.3). The absence of propanediol pathways was supported by the lack of the genus such as *Salmonella* (0% relative abundance in both reactors) coded for gene *bphJ* (Torre, 2002). As for the acrylate pathways, although the gene involved in the intermediate step of converting lactoyl-CoA to acryloyl-CoA was missing (gene *LK433*: mRNA/DNA = 0), the gene *acdA* which participated in converting acryloyl-CoA to propionyl-CoA was still actively expressed. Likely, the acryloyl-CoA pathway was still actively transcribed with acryloyl-CoA generated from amino acid degradation instead of from lactoyl-CoA or lactic acid (Torre, 2002). Two pathways were detected to convert the produced propionyl-CoA to PA: (1) propionyl-P to PA (gene *ackA*); (2) propionyl-adenylate to PA (gene *acs*), and the mRNA/DNA RPKM ratio of genes involved in these two pathways didn’t exhibit any difference (*p* all > .05). In summary, although certain intermediated steps had no gene expression (mRNA/DNA = 0), all other expressed genes involved in PA production didn’t demonstrate any difference in terms of the mRNA/DNA ratio. This similarity in PA production activities and the number of expressed pathways corresponded to the unchanged PA fractions within the two reactors.

**Fig. 3.**
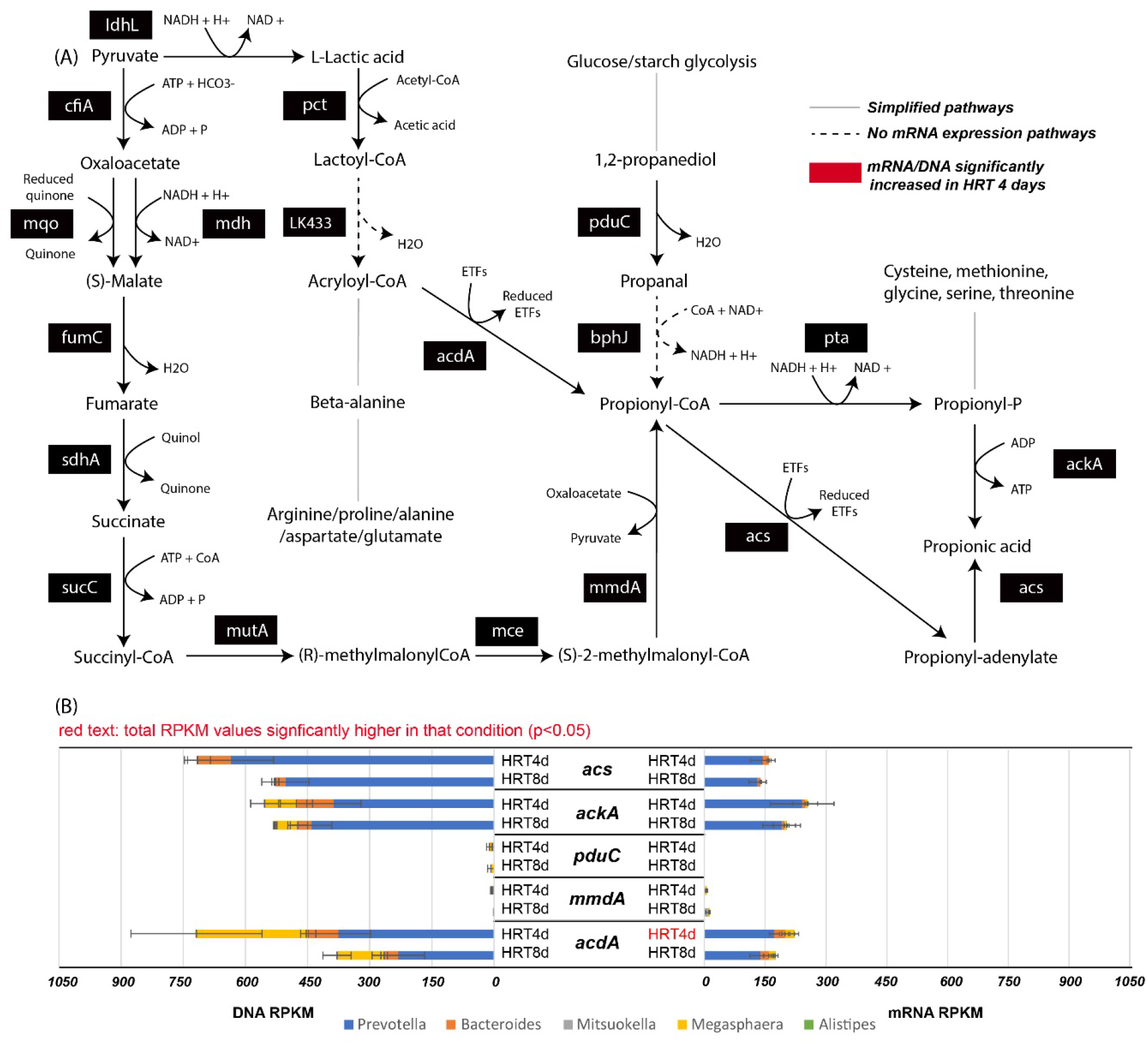
Propionic acid (PA) production metabolic networks and taxonomic origins

It was found that although different genera produced propionyl-CoA via various pathways, no difference was detected between the two HRTs regarding the taxonomic origins in each pathway. *Prevotella, Bacteroides*, and *Megasphaera* were observed to code and express the gene *acdA* involved in the acryloyl-CoA pathway of propionyl-CoA production (HRT 4 days DNA RPKM: 718.29±253.63, HRT 8 days DNA RPKM: 378.89±105.46, *p* = .13; HRT 4 days mRNA RPKM: 221.51±31.47, HRT 8 days mRNA RPKM: 174.87±44.96, *p* = .22). On the other side, the methylmalonyl-CoA pathway of propionyl-CoA production (gene *mmdA*) was mainly encoded and expressed in *Mitsuokella, Megasphaera, Synergistes*, and *Succiniclasticum* (HRT 4 days DNA PRKM:9.16± 7.97, HRT 8 days DNA RPKM: 1.47±3.11, *p* = .23; HRT 4 days mRNA RPKM: 52.49±32.96, HRT 8 days mRNA RPKM: 33.00±23.79, *p* = .46). As for the pathways of converting propionyl-CoA to PA, *Prevotella* was the major genus coded with highest DNA RPKM and expressed with highest mRNA RPKM of genes *acs* and *ackA* involved in the two final steps of PA production (Fig.3), and there is no difference between the two HRTs (all *p* > 0.05). When recalling the structural results, the abundance of *Prevotella* was positively correlated to HRT 8 days. However, through the analysis of metabolic networks, it was found that under either HRTs, *Prevotella* participated in the PA production by converting acryloyl-CoA to propionyl-CoA, and further converting propionyl-CoA to PA through propionyl-P or propionyl-adenylate. These results suggested again that instead of structural changes, the underlying metabolic potential and activities of PA production could be better explained by correspondence to the formed PA production performance.

#### 3.4.3. Butyric acid production

Recalling the performance results, the BA fractions also didn’t exhibit any difference between the two reactors. Within the BA metabolic networks, crotonyl-CoA/butyryl-CoA served as the most important intermediates. Four different pathways were reported for crotonyl-CoA synthesis (Fig.4): (1) acetyl-CoA chain-elongation (gene *crt*); (2) lysine degradation (gene *kal*); (3) glutarate degradation (gene *gcdA*); (4) 4-aminobutyrate pathways (gene *AbfD*). All four pathways converged at producing crotonyl-CoA and followed by the essential step to convert crotonyl-CoA to butyryl-CoA by butyryl-CoA dehydrogenase electron-transferring flavoprotein complex (*Bcd-Etf αβ*)(Vital et al., 2014). This important enzyme complex was documented as EC: 1.3.1.109 and was found in *Megasphaera* (Chowdhury et al., 2015). Although *Megasphaera* was detected as one of the five most abundant genera, no mRNA or DNA was detected for the gene encoded for this complex. However, partial evidence of the enzyme complex as the butyryl-CoA dehydrogenase enzyme (EC: 1.3.8.1) was detected with the expression of gene *acdA/Bcd*. This enzyme also participated in the conversion of acryloyl-CoA to propionyl-CoA. The reasons for the missed detection of *Bcd-Etfαβ* complex were unknown, but it was assumed that crotonyl-CoA was converted to butyryl-CoA via EC: 1.3.8.1 with electron-transfer flavoprotein dimer or other electron bifurcation proteins. Among these four pathways to generate crotonyl-CoA/butyryl-CoA, three of them (acetyl-CoA chain-elongation, lysine degradation, and protein degradation pathways) were detected with all involved genes showing the mRNA/DNA > 0 (no dashed line within the pathways, Fig.4). Gene *gcdA* in the lysine degradation pathway exhibited significantly higher mRNA/DNA ratio under HRT 4 days (*p* = .046). This gene encoded for enzyme glutaconyl-CoA decarboxylase, which was found in *Acidaminococcus*, and acted as a sodium pump (Buckel, 2001). Although this gene exhibited an enhanced mRNA/DNA ratio, all other genes involved in crotonyl-CoA production pathways didn’t demonstrate any changes at the two HRTs. Besides, the genes involved in the last step of BA production (*atoD* and *buk2*) didn’t present any difference in the mRNA/DNA ratio between the two HRTs (*p* = .36 and .21, respectively) which corresponded to the unchanged BA production fractions (*p* = .60). It is suggested that the expression of the terminal step might be more correlated with the performance than the expression of intermediate steps, which could be considered the indicators of the performance.

**Fig. 4.**
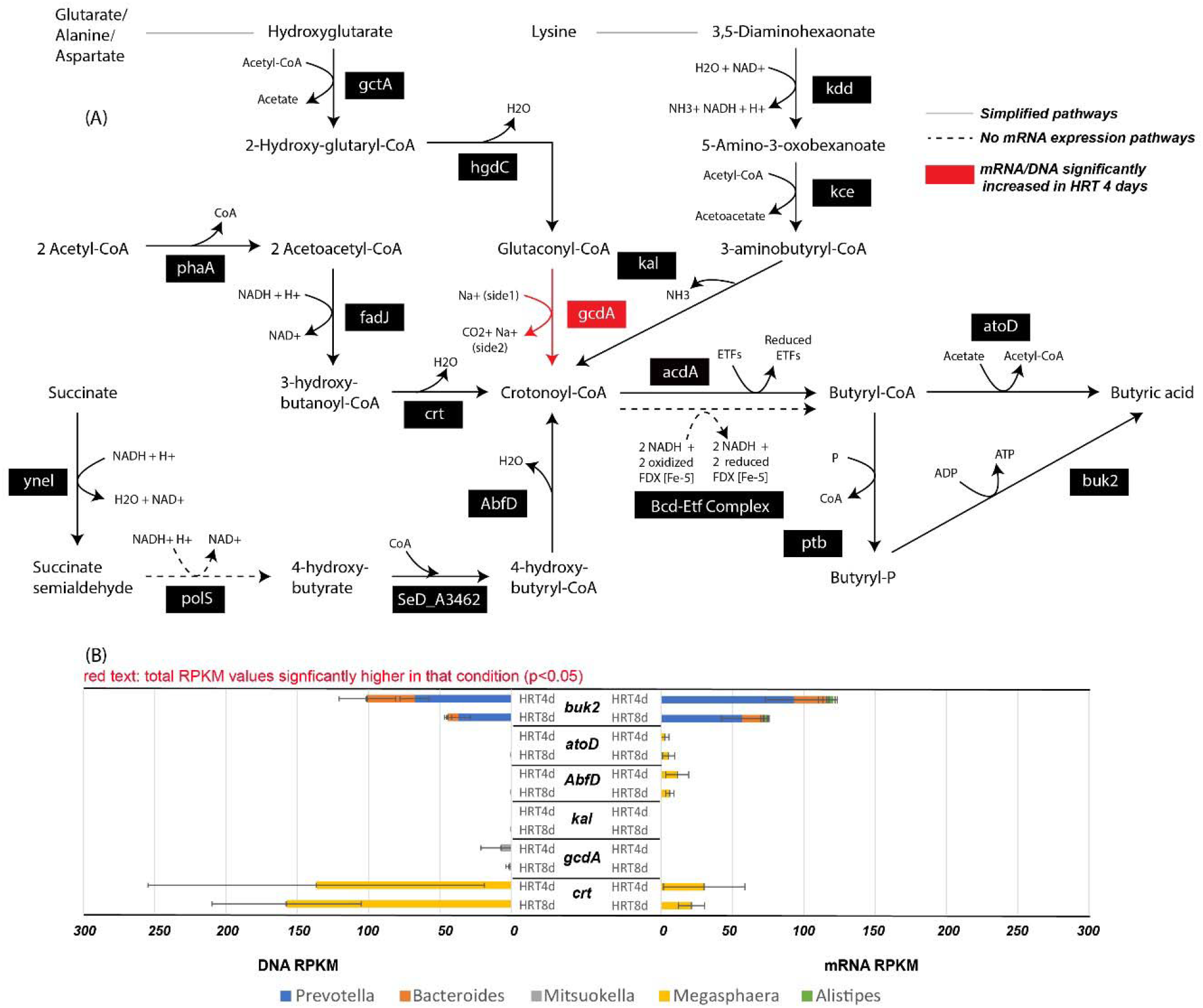
Butyric acid (BA) production metabolic networks and taxonomic origins

When exploring the taxonomic origins of each BA production pathway, a potential cross-feeding was found between the top five abundant groups. It was noticed that *Megasphaera* was the only genus found with the DNA RPKM (HRT 4 days: 136.82±117.92, HRT 8 days: 157.61±52.28, *p* = .80, Fig.4) and mRNA RPKM (HRT 4 days: 30.18±28.52, HRT 8 days: 21.29±9.08, *p* = .65, Fig.4) of gene *crt*. However, other abundant genera including *Prevotella, Bacteroides*, and *Alistipes* were only found with the DNA RPKM (HRT 4d: 48.01±30.59, HRT 8d 44.74±11.13, *p* = .87) and mRNA PRKM (HRT 4d: 120.64±29.20, HRT 8d: 75.18±17.50, *p* = .10) of gene *buk2*. It is possible that *Megasphaera* elongated the acyl-chain and generated the essential intermediates crotonyl-CoA/butyryl-CoA, while *Prevotella, Bacteroides*, and *Alistipes* directly utilized the formed crotonyl-CoA/butyryl-CoA and further converted it to BA. This result indicated a complex substrate partial-degradation and transfer process within the acidification communities. The production of the end products might be jointly completed by the cross-feeding between distinct genera generating the intermediates and transferring them to other groups as the substrate to form the final targeted metabolites.

#### 3.4.4. Valeric acid production

VA was the only increased acid under HRT 8 days (*p* = .035) and several studies also reported that when increasing the HRT, the acid chain was elongated with ethanol and lactate being utilized as the electron donors (Contreras-Davila et al., 2020; De Groof et al., 2021; Wu et al., 2018). However, no specific evidence could be found from the analysis of metabolic networks to confirm the enhanced expression of chain-elongation genes under longer HRT. The three detected genes *fadA, fadJ*, and *crt* didn’t exhibit any difference in terms of mRNA/DNA ratio (*p* = .50, .12, .66, respectively). As for the taxonomic origins, *Megasphaera* was mainly responsible for the chain-elongation process, which exhibited the highest DNA RPKM (HRT 4 days: 136.82±117.92, HRT 8 days: 157.61±52.28, *p* = 0.80, Fig.5) and mRNA RPKM (HRT 4 days: 30.18±28.52, HRT 8 days: 21.29±9.08, *p* = .65, Fig.5) of gene *crt. Bacteroides* also contributed to the chain-elongation process by expressing gene *fadJ* (HRT 4 days mRNA RPKM: 9.90±4.03, HRT 8 days: 3.73±1.04, *p* = .11).

**Fig. 5.**
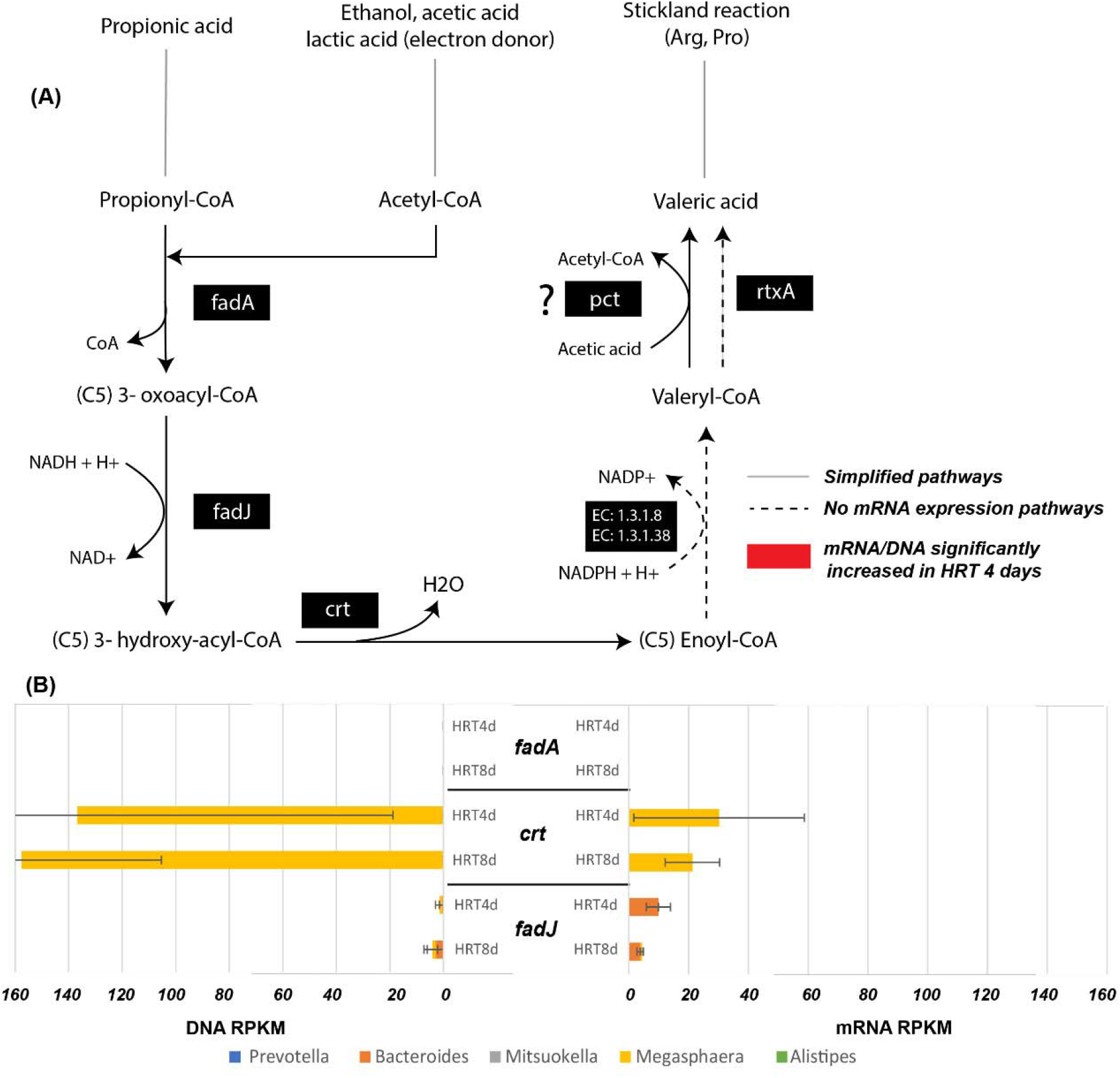
Valeric acid (VA) production metabolic networks and taxonomic origins

Besides, genes encoded for enzyme EC: 1.3.1.8 or EC: 1.3.1.38 and gene *rtxA* were not found in both reactors (mRNA/DNA RPKM = 0, Fig.5 dash lines). The production of VA without detecting continuous expression of genes involved in the chain-elongation pathways suggested that there might exist other unknown pathways or enzymes, which catalyzed the reactions during the reverse beta-oxidation. For example, VA was also reported to be produced from protein degradation. This pathway was found in proteolytic species such as *Clostridium*, which generated VA via reductive deamination of single amino acids, or by an oxidation-reduction reaction between paired amino acids known as the Stickland reaction (Neumann-Schaal et al., 2019). The genus *Clostridium* was presented in both reactors with a relative abundance of 0.93 ± 0.07% (HRT 4 days) and 1.25±0.36 % (HRT 8 days), respectively. Although the abundance didn’t show any difference between the two reactors (*p* = .26), likely, the potential or extant functions in VA production from the Stickland reaction were enhanced under HRT 8 days. More metabolic networks within different protein degradation to VA production could be further explored in the future.

## 4. Conclusions

Although distinct microbial structures were shaped by the two selected HRTs (4 days and 8 days), the total VFA yield and dominantly produced propionic acid (PA) were unchanged. The meta-omics results revealed similar potential (DNA RPKM) and extant (mRNA RPKM) functions in the customized acidification metabolic networks. Compared to other acids, PA production exhibited the highest potential and extant functions in both reactors, and the meta-omics analysis indicated that the most abundant genus *Prevotella* produced PA mainly through the acryloyl-CoA pathway. Based on our results arrested AD processes can achieve stable and similar VFA production performance under different HRTs, due to the convergence and redundancy in microbial acidification functionality despite divergence in microbial structure.

## Supporting information

https://docs.google.com/document/d/1nGSzclcJ33xX8GFobrTSRcjIQE75k6p6/edit?usp=sharing&ouid=100587333082155714262&rtpof=true&sd=true

## Acknowledgments

This work was sponsored by the Water Research Foundation with the project number WRF 4900.

## Appendix A.

**supplementary data**

---

**Fig.1.** (A): Temporal change of the relative abundance of bacteria (only genera with relative abundance > 1%) and all archaea communities. (B): Type II scaled triplot RDA of the microbial ecology associated with different HRTs and food waste properties. Samples were displayed in circles (HRT 4 days) and squares (HRT 8 days). All genera were indicated by red arrows. Explanatory factors (HRTs and food waste properties) were presented by blue arrows. The significance of different explanatory factors was summarized in the table (*P* < 0.05 were marked with **).

**Fig.2.** (A): AA production metabolic networks: The pathways with mRNA/DNA>0 were regarded as expressed and presented in solid lines. Pathways with mRNA/DNA=0 were regarded as having no expression and presented in dashed lines. Simplified pathways were presented in the grey line. The significantly increased mRNA/DNA of genes under HRT 4 days were highlighted in red. (B): The taxonomic origins of AA production pathways: Contributions from the top 5 most abundant genera to maker genes’ DNA and mRNA RPKM. Maker genes were the genes involved in the final step of each acid production pathway. The significantly changed DNA or mRNA RPKM between the two HRTs were highlighted in red text.

**Fig.3.** (A): PA production metabolic networks. (B) Taxonomic origins of each PA production pathway

**Fig.4.** (A) BA production metabolic networks. (B) Taxonomic origins of each BA production pathway

**Fig.5.** (A) VA production metabolic networks. (B) Taxonomic origins of each VA production pathway

