## Supplementary material for "Divergent microbial structure still results in convergent microbial function during arrested anaerobic digestion of food waste at different hydraulic retention times": https://docs.google.com/document/d/1nGSzclcJ33xX8GFobrTSRcjIQE75k6p6/edit?usp=sharing&ouid=100587333082155714262&rtpof=true&sd=true

**Equations:**

$$VFA yield\left( \% \right)=\frac{Eff COD\left( \mathrm{VFA} \right)-Inf COD \left( \mathrm{VFA} \right)}{Inf tCOD}\times100\% \left( 1 \right)$$

$$Methane yield (\%) =\frac{Total COD \left( CH_{4} \right)}{Inf tCOD} \times100\% \left( 2 \right)$$

$$Individual acid fraction of total produced VFA (\%) =\frac{individual acid concentration \times COD equivalent conversion factor}{Sum of four acids COD equivalent}\times100\% \left( 3 \right)$$

Global function DNA reads-based percentage

$=\frac{number of mapped DNA reads per global functional classification}{total number of DNA reads per sample}$	(4)

Global function mRNA reads-based percentage

$=\frac{number of mapped mRNA reads per global functional classification}{total number of mRNA reads per sample}$	(5)

DNA RPKM $=\frac{number of mapped DNA reads per gene \times{10}^{6}}{\frac{gene length}{1kb}\times total number of DNA reads per sample}$	(6)

mRNA RPKM $=\frac{number of mapped mRNA reads per gene \times{10}^{6}}{\frac{gene length}{1kb}\times total number of mRNA reads per sample}$	(7)

Absolute transcriptional levels$=\frac{RPKM DNA}{RPKM mRNA}$	(8)

Taxonomy contribution DNA RPKM = $\frac{number of mapped DNA reads per genus per gene}{number of mapped DNA reads per gene}\times RPKM DNA per gene$	(9)

Taxonomy contribution to mRNA RPKM = $\frac{number of mapped mRNA reads per genus per gene}{number of mapped mRNA reads per gene}\times RPKM mRNA per gene$	(10)

The methane gas was only measured during the operational day 299 to 422. The COD conversion factors were used as followed: 4g COD/g CH4, 1.07 g COD/g acetate, 1.51g COD /g propionate, 1.82g COD/ g butyrate, and 2.04 g COD/g valerate. All values were calculated after both reactors reached a relatively stable performance (the effluent VFA concentration had less than 5% relative standard deviation (RSD)).

**Tables and Figures:**

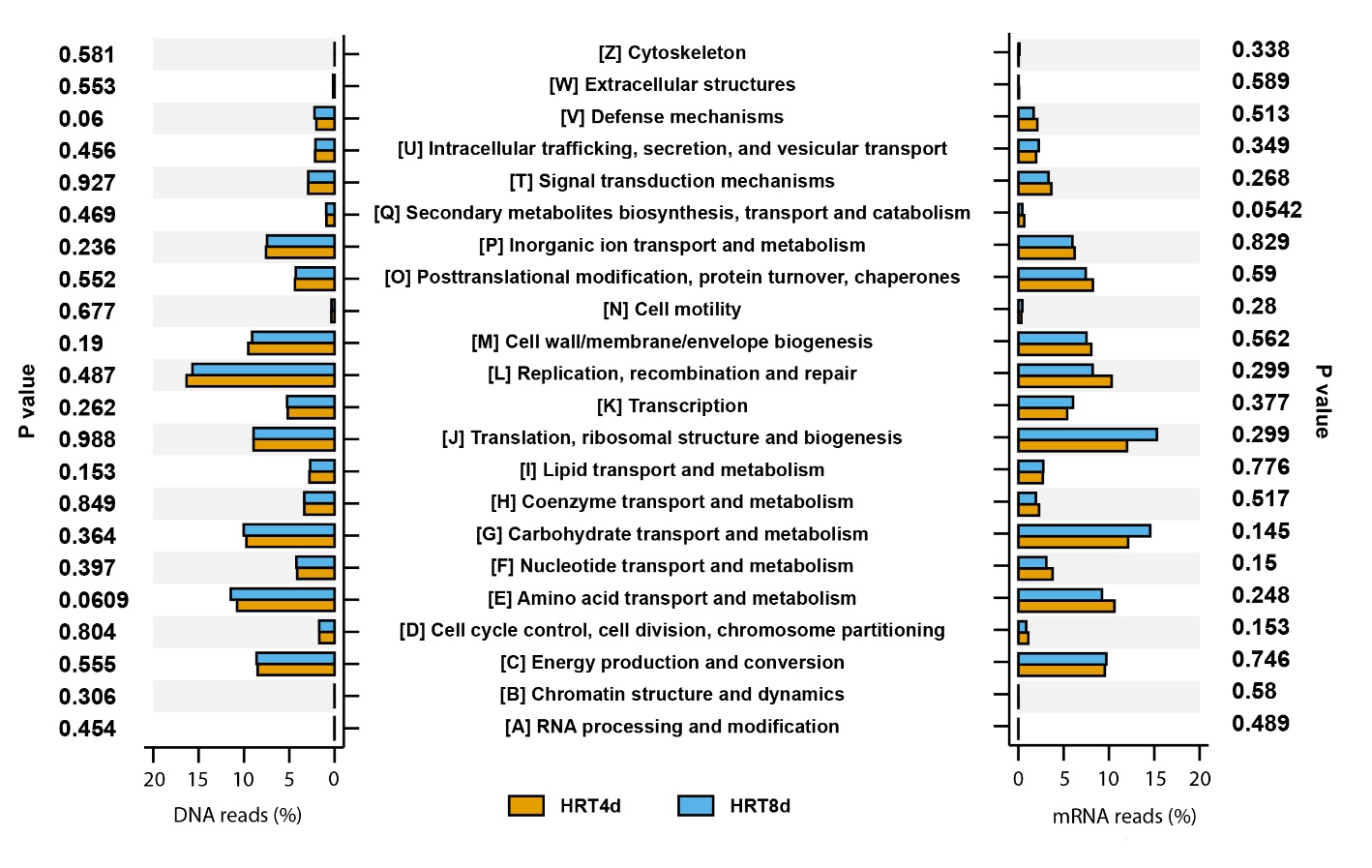
.

**Supplement Fig.1.** Averaged reads-based percentage (mapped reads/total reads %) (%) of genes annotated to the eggNOG database

**Supplement Table 1**. Influent Feedstock Characteristics

| Operatio-nal days | sCOD/ tCOD  Ratio | Total Protein  (g BSA/L) | Total lipid  (%weight /influent) | Total carbohydrate  (g glucose/L) | TS  (g/L) | VS  (g/L) | Soluble protein  (g BSA/L) | Soluble carbohydrate  (g glucose/L) |
| --- | --- | --- | --- | --- | --- | --- | --- | --- |
| 55-83 | 35.72$\pm8.42$ | 2.55$\pm0.25$ | 1.13$\pm0.06$ | 6.70$\pm0.79$ | 33.22$\pm0.80$ | 31.31$\pm0.81$ | 1.48$\pm0.03$ | 2.69$\pm0.59$ |
| 83-114 | 33.93$\pm5.29$ | 2.51$\pm0.25$ | 1.18$\pm0.16$ | 5.15$\pm1.65$ | 35.13$\pm1.47$ | 32.90$\pm1.47$ | 1.49$\pm0.05$ | 1.89$\pm0.21$ |
| 114-144 | 5.38$\pm1.74$ | 1.78$\pm0.08$ | 0.91$\pm0.01$ | 2.29$\pm0.33$ | 31.74$\pm6.27$ | 30.45$\pm5.86$ | 1.33$\pm0.07$ | 0.06$\pm0.02$ |
| 144-175 | 10.36$\pm0.71$ | 2.31$\pm0.04$ | 17.89$\pm0.56$ | 13.05$\pm0.00$ | 23.17$\pm0.36$ | 22.24$\pm0.40$ | 1.35$\pm0.07$ | 0.01$\pm0.01$ |
| 175-205 | 36.05$\pm3.82$ | 4.18$\pm0.12$ | 0.70$\pm0.03$ | 31.39$\pm4.39$ | 42.58$\pm3.79$ | 41.50$\pm3.64$ | 1.84$\pm0.02$ | 5.67$\pm2.25$ |
| 205-236 | 33.35$\pm3.93$ | 6.21$\pm0.91$ | 1.01$\pm0.02$ | 32.38$\pm5.12$ | 60.14$\pm5.45$ | 58.26$\pm5.40$ | 2.57$\pm0.11$ | 2.41$\pm0.20$ |
| 236-267 | 71.80$\pm7.22$ | 6.06$\pm0.70$ | 0.78$\pm0.04$ | 6.49$\pm1.73$ | 27.46$\pm1.94$ | 25.25$\pm1.77$ | 2.95$\pm0.11$ | 1.36$\pm0.07$ |
| 267-297 | 27.58$\pm9.07$ | 7.03$\pm0.10$ | 2.66$\pm1.68$ | 27.41$\pm4.39$ | 55.34$\pm2.29$ | 52.66$\pm2.30$ | 3.33$\pm0.21$ | 2.92$\pm0.35$ |
| 297-328 | 22.62$\pm5.97$ | 5.54$\pm1.07$ | 2.16$\pm0.90$ | 18.02$\pm3.36$ | 38.53$\pm2.26$ | 36.33$\pm2.35$ | 2.32$\pm0.13$ | 2.02$\pm0.26$ |
| 328-358 | 17.65$\pm2.87$ | 4.05$\pm2.05$ | 1.65$\pm0.12$ | 8.63$\pm2.33$ | 21.72$\pm2.24$ | 20.01$\pm2.41$ | 1.30$\pm0.05$ | 1.12$\pm0.18$ |
| 358-388 | 13.99$\pm0.70$ | 3.70$\pm2.14$ | 0.87$\pm0.02$ | 6.59$\pm0.34$ | 39.16$\pm15.41$ | 36.60$\pm13.75$ | 1.57$\pm0.12$ | 1.27$\pm0.02$ |
| 388-422 | 31.71$\pm11.80$ | 2.11$\pm0.06$ | 0.37$\pm0.02$ | 4.61$\pm0.24$ | 31.12$\pm0.30$ | 28.81$\pm0.27$ | 1.44$\pm0.02$ | 0.89$\pm0.01$ |

Note: All results were measured by duplicates.

**Supplement Table 2.** All genes and corresponding encoded enzymes involved in the acidification process and methanogenesis process (three pathways)

| Category | EC number | gene name | gene length | encoded enzymes |
| --- | --- | --- | --- | --- |
| FA production | EC: 2.3.1.54 | *pflB* | 2244 | 2.3.1.54 Formate C-acetyltransferase |
| LA production | EC: 1.1.1.27 | *ldhL* | 933 | 1.1.1.27 L-lactate dehydrogenase |
| AA production | EC: 1.2.7.1 | *nifj* | 3547 | 1.2.7.1 Pyruvate synthase |
|  | EC: 2.8.3.1 | *pct* | 1554 | 2.8.3.1 Propionate CoA-transferase |
|  | EC: 1.2.7.1 | *nifj* | 3547 | 1.2.7.1 Pyruvate synthase |
|  | EC: 2.3.1.8 | *pta* | 1027 | 2.3.1.8 Phosphate acetyltransferase |
|  | EC: 2.7.2.1 | *ackA* | 1205 | 2.7.2.1 Acetate kinase |
|  | EC:1.2.5.1 | *poxB* | 1764 | 1.2.5.1 Pyruvate dehydrogenase (quinone) |
|  | EC: 1.2.3.3 | *spxB* | 1735.5 | 1.2.3.3 Pyruvate oxidase |
|  | EC:3.6.1.7 | *yccX* | 254 | 3.6.1.7 Acylphosphatase |
|  | EC: 2.7.2.1 | *ackA* | 1205 | 2.7.2.1 Acetate kinase |
|  | EC: 1.2.7.1 | *nifj* | 3547 | 1.2.7.1 Pyruvate synthase |
|  | EC: 6.2.1.1 | *acs* | 1959 | 6.2.1.1 Acetate--CoA ligase |
|  | EC:6.2.1.13 | *pauA* | 2061 | 6.2.1.13 Acetate--CoA ligase (ADP-forming) |
|  | EC: 1.2.7.1 | *nifj* | 3547 | 1.2.7.1 Pyruvate synthase |
|  | EC: 1.2.1.10 | *mhpF* | 951 | Acetaldehyde dehydrogenase (acetylating) |
|  | EC1.2.1.3 | *aldH/putA1* | 1492.2 | 1.2.1.3 Aldehyde dehydrogenase (NAD(+)) |
| PA production | EC:2.3.1.9 | *phaA/fadA/atoB/thlA/yqeF* | 1178 | 2.3.1.9 Acetyl-CoA C-acetyltransferase |
|  | EC: 1.1.1.27 | *ldhL* | 933 | 1.1.1.27 L-lactate dehydrogenase |
|  | EC: 2.8.3.1 | *pct* | 1554 | 2.8.3.1 Propionate CoA-transferase |
|  | EC: 4.2.1.54 | *LK433_RS01340* | 1206 | Lactoyl-CoA dehydratase |
|  | EC: 1.3.8.1 | *acdA/bcd/ACADS* | 1518 | 1.3.8.1 Short-chain acyl-CoA dehydrogenase |
|  | EC: 2.3.1.8 | *pta* | 1027 | 2.3.1.8 Phosphate acetyltransferase |
|  | EC: 2.7.2.1 | *ackA* | 1205 | 2.7.2.1 Acetate kinase |
|  | EC: 6.2.1.1 | *acs* | 1959 | 6.2.1.1 Acetate--CoA ligase |
|  | EC: 6.4.1.1 | *cfiA_2/cfiA/pyc* | 2220 | 6.4.1.1 Pyruvate carboxylase |
|  | EC: 1.1.1.37 | *mdh* | 939 | Malate dehydrogenase |
|  | EC: 1.1.5.4 | *mqo* | 1647 | Malate dehydrogenase (quinone) |
|  | EC: 4.2.1.2 | *fumC* | 843 | Fumarate hydratase |
|  | EC: 1.3.5.4 | *sdhA/frdA/frdB* | 1669.2 | 1.3.5.4 Fumarate reductase (quinol) |
|  | EC: 6.2.1.5 | *sucC/sucD* | 968.1429 | 6.2.1.5 Succinate--CoA ligase (ADP-forming) |
|  | EC: 2.8.3.18 | *aarC* | 1500 | 2.8.3.18 Succinyl-CoA: acetate CoA-transferase |
|  | EC: 5.4.99.2 | *mutA/scpA* | 2049 | 5.4.99.2 Methylmalonyl-CoA mutase |
|  | EC:5.1.99.1 | *mce* | 414 | 5.1.99.1 Methylmalonyl-CoA epimerase |
|  | EC: 2.1.3.1 | *mmdA* | 1534.5 | methylmalonyl-CoA carboxytransferase; |
|  | EC: 4.2.1.28 | *pduC* | 1602 | Propanediol dehydratase |
|  | EC: 1.2.1.87 | *bphJ* | 1197 | Propanal dehydrogenase (CoA-propanoylating) |
| BA production | EC:2.3.1.9 | *phaA/fadA/atoB/thlA/yqeF* | 1178 | 2.3.1.9 Acetyl-CoA C-acetyltransferase |
|  | EC: 1.1.1.35 | *fadJ/fadB/yfcX* | 975 | 1.1.1.35 3-hydroxyacyl-CoA dehydrogenase |
|  | EC:4.2.1.17 | *crt2/fadJ/fadB/echA15/paaF* | 783 | 4.2.1.17 Enoyl-CoA hydratase |
|  | EC: 1.3.1.44/1.3.1.86 | *fabV* | 1197 |  |
|  | EC: 2.8.3.8 | *atoD/atoA/ctfA/ctfB* | 660 | 2.8.3.8 acetate CoA/acetoacetate CoA-transferase beta subunit |
|  | EC: 2.3.1.19 | *ptb* | 1065 | 2.3.1.19 Phosphate butyryltransferase |
|  | EC:2.7.2.7 | *buk2* | 1062 | 2.7.2.7 Butyrate kinase |
|  | EC: 1.1.1.16 | *polS/AtuSorbD* | 774 | Galactitol 2-dehydrogenase |
|  | EC: 2.8.3.12 | *gctA/gctB* | 864 | Glutaconate CoA-transferase |
|  | EC: 7.2.4.5 | *gcdA* | 1242 | Glutaconyl-CoA decarboxylase |
|  | EC: 1.4.1.11 | *kdd* | 1059 | L-erythro-3,5-diaminohexanoate dehydrogenase |
|  | EC: 1.2.1.24 | *yneI* | 1389 |  |
|  | EC: 2.3.1.247 | *kce* | 813 | 3-keto-5-aminohexanoate cleavage enzyme |
|  | EC: 4.3.1.14 | *kal* | 390 | 3-aminobutyryl-CoA ammonia-lyase |
| VA production | EC: 2.3.1.16 | *FadA/atoB* | 1164 | Acetyl-CoA C-acyltransferase |
|  | EC: 1.1.1.35 | *fadJ* | 975 | 3-hydroxyacyl-CoA dehydrogenase |
|  | EC: 4.2.1.17 | *crt2/crt* | 783 | Enoyl-CoA hydratase |
|  | EC: 1.3.1.38 | *qorA_2* | 981 |  |
|  | EC: 3.1.2.22 | *rtxA* | 825 | Palmitoyl-protein hydrolase |
| Beta-oxidation | EC: 6.2.1.3 | *fadD* | 1487 | 6.2.1.3 Long-chain-fatty-acid--CoA ligase |
| methanogenesis (three pathways) | EC: 1.2.7.12 | *fwdA/fwdB/fwdC/fwdD/fwdH* | 1026 | 1.2.7.12 Formylmethanofuran dehydrogenase |
|  | EC: 2.3.1.101 | *ftr1* | 891 | 2.3.1.101 Formylmethanofuran--tetrahydromethanopterin N-formyltransferase |
|  | EC: 3.5.4.27 | *mch* | 966 | 3.5.4.27 Methenyltetrahydromethanopterin cyclohydrolase |
|  | EC: 1.5.98.1 | *mtd* | 831 | 1.5.98.1 methylenetetrahydromethanopterin dehydrogenase |
|  | EC: 1.12.98.2 | *hmd* | 1017 | 1.12.98.2 5,10-methenyltetrahydromethanopterin hydrogenase |
|  | EC: 2.1.1.86 | *mtrA-H/cmtA* | 555 | 2.1.1.86 Tetrahydromethanopterin S-methyltransferase |
|  | EC: 2.8.4.1 | *mcrA* | 1252 | 2.8.4.1 Coenzyme-B sulfoethylthiotransferase |
|  | EC: 1.8.7.3 | *CoB/CoM / hdrABC/hdrA1B1C1* | 1155 | 1.8.7.3 Ferredoxin:CoB-CoM heterodisulfide reductase |
|  | EC: 2.1.1.250 | *MttB* | 1515 | 2.1.1.250 [Trimethylamine--corrinoid protein] Co-methyltransferase |
|  | EC: 2.1.1.249 | *MtbB* | 1410 | 2.1.1.249 [Dimethylamine--corrinoid protein] Co-methyltransferase |
|  | EC: 2.1.1.248 | *MtmB* | 1380 | 2.1.1.248 [Methylamine--corrinoid protein] Co-methyltransferase |
|  | EC: 2.1.1.246 | *mtbA/mtaA* | 1116 | 2.1.1.246 methyltransferase |

**Supplement Table 3:** Relative abundance % results from whole genome sequencing

| Phylum | Genus | HRT8_1 | HRT8_2 | HRT8_3 | HRT4_1 | HRT4_2 | HRT4_3 |
| --- | --- | --- | --- | --- | --- | --- | --- |
| Bacteroidetes | *Bacteroides* | 12.36 | 9.86 | 7.89 | 14.11 | 24.65 | 15.04 |
| Bacteroidetes | *Prevotella* | 29.75 | 30.98 | 32.57 | 37.80 | 33.44 | 36.95 |
| Bacteroidetes | *Alistipes* | 5.10 | 5.01 | 5.35 | 4.11 | 2.27 | 3.25 |
| Bacteroidetes | *Parabacteroides* | 0.94 | 0.89 | 0.84 | 1.39 | 2.33 | 1.91 |
| Proteobacteria | *Alcaligenes* | 0.29 | 0.32 | 0.22 | 0.68 | 1.13 | 0.23 |
| Proteobacteria | *Succinatimonas* | 5.55 | 3.46 | 2.44 | 1.35 | 0.83 | 2.22 |
| Proteobacteria | *Succinivibrio* | 4.73 | 2.44 | 2.16 | 1.11 | 0.74 | 1.98 |
| Synergistetes | *Pyramidobacter* | 1.64 | 3.21 | 1.22 | 0.59 | 0.50 | 0.70 |
| Synergistetes | *Synergistes* | 3.33 | 3.93 | 3.77 | 2.99 | 2.58 | 3.27 |
| Actinobacteria | *Bifidobacterium* | 1.55 | 3.81 | 2.02 | 1.50 | 1.10 | 2.78 |
| Actinobacteria | *Olsenella* | 3.38 | 3.18 | 3.42 | 2.98 | 2.88 | 3.04 |
| Firmicutes | *Clostridium* | 1.06 | 1.03 | 1.66 | 0.85 | 0.97 | 0.96 |
| Firmicutes | *Eubacterium* | 0.87 | 1.13 | 1.60 | 0.64 | 0.78 | 0.87 |
| Firmicutes | *Butyrivibrio* | 0.64 | 0.76 | 1.18 | 0.44 | 0.52 | 0.68 |
| Firmicutes | *Roseburia* | 1.58 | 0.79 | 0.88 | 0.40 | 0.44 | 0.43 |
| Firmicutes | *Anaerolactibacter* | 1.93 | 2.06 | 2.20 | 1.86 | 1.36 | 1.08 |
| Firmicutes | *Acidaminococcus* | 1.99 | 1.53 | 2.13 | 2.10 | 1.47 | 1.93 |
| Firmicutes | *Mitsuokella* | 6.16 | 5.68 | 5.61 | 4.23 | 4.48 | 3.52 |
| Firmicutes | *Selenomonas* | 4.63 | 3.46 | 6.04 | 4.11 | 1.65 | 3.04 |
| Firmicutes | *Megasphaera* | 3.69 | 3.18 | 4.07 | 8.14 | 4.44 | 5.75 |
| Archaea | *Candidatus Methanomethylophilus* | 1.34 | 3.11 | 1.31 | 1.38 | 1.06 | 1.13 |
| Archaea | *Methanobrevibacter* | 0.52 | 0.92 | 0.92 | 0.49 | 0.42 | 0.59 |
| Others | *Others* | 6.97 | 9.25 | 10.50 | 6.74 | 9.95 | 8.65 |
